# From cells to organism – how natural selection causes metabolic scaling

**DOI:** 10.1101/2025.07.24.666547

**Authors:** Peter F. Pelz

**Author notes:** Contributing authors.

## Abstract

The cell is the power station of life. Surprisingly, to date there is still no metabolic scaling theory that links cellular respiration to organismal metabolism and predicts the ‘mouse-to-elephant curve’, also known as Kleiber’s law, in an approach that is consistent with physicochemical principles. This paper shows that for a consistent model, the novel concept of the optimised Metabolic Module (MM) is the missing link between cell and organism. It is shown how evolutionary selection under resource scarcity optimises the MM towards (a) lightweight design and (b) resource efficiency. Thus, Darwin’s evolution by natural selection is simulated by model-based optimisation. The final general model presented is complete (for the entire mass range of the organism of different taxonomic classes), concise (it uses only five scale-invariant physicochemical constants), clear (it predicts all metabolic rates within the uncertainty range of a scale model observed in measurements) and consistent with Murray’s law of capillary blood flow and cell metabolism. The model features observed asymptotes for both ‘small’ protists and ‘large’ endotherms. It predicts the mass-dependent metabolic rate of protists, planarians, ectotherms and endotherms with the usual uncertainty of any scaling theory. It finally turns out that Kleiber’s law is an asymptote of the derived general model, namely for the case of diffusion-limited cell metabolism.

## 1 Introduction

### Cellular architecture of life

The idea of a cellular architecture of life is common. In this concept the growth of the organism is achieved by the multiplication of scale-invariant cells, and the metabolic rate of the organism results from this multiplication of cells that are regarded as metabolic power stations. To account for the chemical nature of cellular respiration, the chemical rate constant *k*, measured in ‘per time’ units, is the most relevant physico-chemical parameter. *k* is the rate of the slowest reaction step in the multi-step reaction within cells.

In ‘small’ organisms, the metabolic rate is proportional to *k* and to the body mass *m* of the organism. This is to be expected in a complete cellular architecture of life. However, the situation is different for ‘large’ organisms: their metabolism is proportional to *k*^3/8^ and *m*^3/4^, as will be shown in this paper.

‘Small’ organisms are protists such as plankton. ‘Large’ organisms include ectotherms (cold-blooded organisms) such as reptiles and amphibians and endotherms (warm-blooded organisms) such as birds and mammals and most planarians (flat-worms) as their typical size is above the diffusion length.

As will be shown, the diffusion length is the relevant measure for distinguishing between ‘large’ and ‘small’, a physiological parameter determined by the diffusion coefficient and the rate constant *k* of cell metabolism.

### Metabolic rate of ‘small’ organisms

In the following, we will first look at the metabolism of the ‘small’ organism. The resulting scaling equation is well known. Nevertheless, the derivation is valuable because some of the arguments are used below to derive a general metabolic scaling theory that is valid for ‘small’ and ‘large’ organisms. As a reminder, oxygen molecules are transported through the cell tissue by diffusion, resulting from a concentration field *c* limited by the available oxygen concentration: *c* ≤ *c*_avail_.

In compliance with Fick’s first law of diffusion, the transport speed of the molecules is proportional to their molecular diffusion coefficient 𝒟 in the cell tissue multiplied by the gradient of the concentration field |Δ |*c < c*_avail_*/δ*. The longest transport path for molecules is equal to the thickness *δ* of the cell tissue. Simple organisms such as plankton have a tissue thickness that is much smaller than the diffusion length, *δ*_diff_ := (𝒟*/k*)^1*/*2^ ≫*δ*. In these ‘small’ organisms, the diffusion time ∼*δ*^2^*/𝒟* of oxygen through the cell tissue is much shorter than the reaction time 1*/k* in the cells.

In the most elementary model of reaction kinetics, the reaction rate, i.e. the oxygen consumption per unit time and unit volume of cell tissue *ċ*, is proportional to the rate constant *k* and the oxygen concentration −*ċ* = *k c*_avail_.

Therefore, the molar flow rate *J* of oxygen molecules for ‘small’ organisms with a volume *V* is determined entirely by the reaction rate of glucose oxidation: *J* = *J*_react_ = − *∫*_*V*_ *ċ* d*V* = *k c*_avail_*V*.

With the abbreviation *Q* := *J/c*_avail_ the kinematic character of this mass balance becomes obvious, i.e. *Q* = *k V*. With the energy density *e* := *E/V* and the mass density *ϱ* := *m/V* of the cell tissue, the metabolic rate, measured as heat production rate, can be derived using the transformations *Ė* = *e Q* and *m* = *ϱ V*

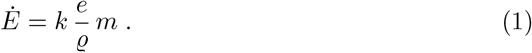

The energy density *e* of the cell tissue is given by the product of the oxygen concentration and the molar free Gibbs energy for the exothermic oxidation of nutrients within the cell, i.e. *e* = *c*_avail_ |Δ*G*^*°*^|.

The given derivation of the scaling equation *Ė*∝ *m* is consistent with the idea of a cellular architecture of life: the cell is the power station of life. Equation (1) is therefore nothing more than a summation of the power contributions of all the cells of a small organism.

### Diffusion limit on growth

The idea of a cellular architecture of life has a limitation: for an organism with a mass *m* far above the diffusion limit on growth, i.e. 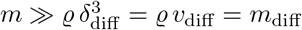, the diffusive supply with oxygen is insufficient to prevent cell death. As shown in this paper, with increasing mass of the organism, this leads to a change from the asymptote *Ė* ∝ *m* for *m ≪ m*_diff_ to the asymptote *Ė ∝ m*^3/4^ for *m ≫ m*_diff_. The latter asymptote is known as Kleiber’s law [1] or as the ‘mouse-to-elephant curve’, see figs. 2 and 3. This asymptote is explained in this paper by a diffusion-limited reaction, in which the diffusion flow is maximised by an optimal modular design.

To date, there is (i) no general metabolic theory that applies to all organisms, from a plankton with a mass of only 0.01 picogram to a blue whale with a mass of more than 100 tonnes. This general theory must include both asymptotes mentioned above; there is (ii) no consistent theory linking cell metabolism to organismal metabolism; and there is (iii) no theory that is consistent with the physiology of blood flow in capillaries [2–4].

The theory presented here fills this gap. It links cell metabolism and the metabolism of the organism via the Metabolic Module (MM), which has been optimised to the size of the animal through natural selection. The concept of a scalable MM is presented here for the first time. The results might be useful for model-based drugs and tissue design.

Darwin’s evolution by natural selection is simulated as model-based optimisation. The solution of the optimisation contributes to the understanding of the metabolism of organisms; the scaling of the metabolic rate resulting from evolution under resource scarcity is explained axiomatically. This new approach is discussed in the following.

## 2 Modular architecture of life

The new idea of an optimised modular architecture of life goes beyond the purely cellular image, as the theory, its asymptotes and its validation will show below.

Every organ and every muscle is supplied with nutrients and oxygen via a capillary network or, to describe the topology more specifically, via a capillary bundle [5] consisting of a large number of modules, see fig. 1. Others [2, 3] consider the capillaries of these modules to be scale-invariant like cells, which is in contradiction to measurements [6, 7] and conflicts with Murray’s law of capillary flow [8]. In contrast, the concept of an optimised Metabolic Module (MM) resolves this validation problem and the consistency conflict of existing metabolic scaling theories [2, 3], as shown below.

**Fig. 1.**
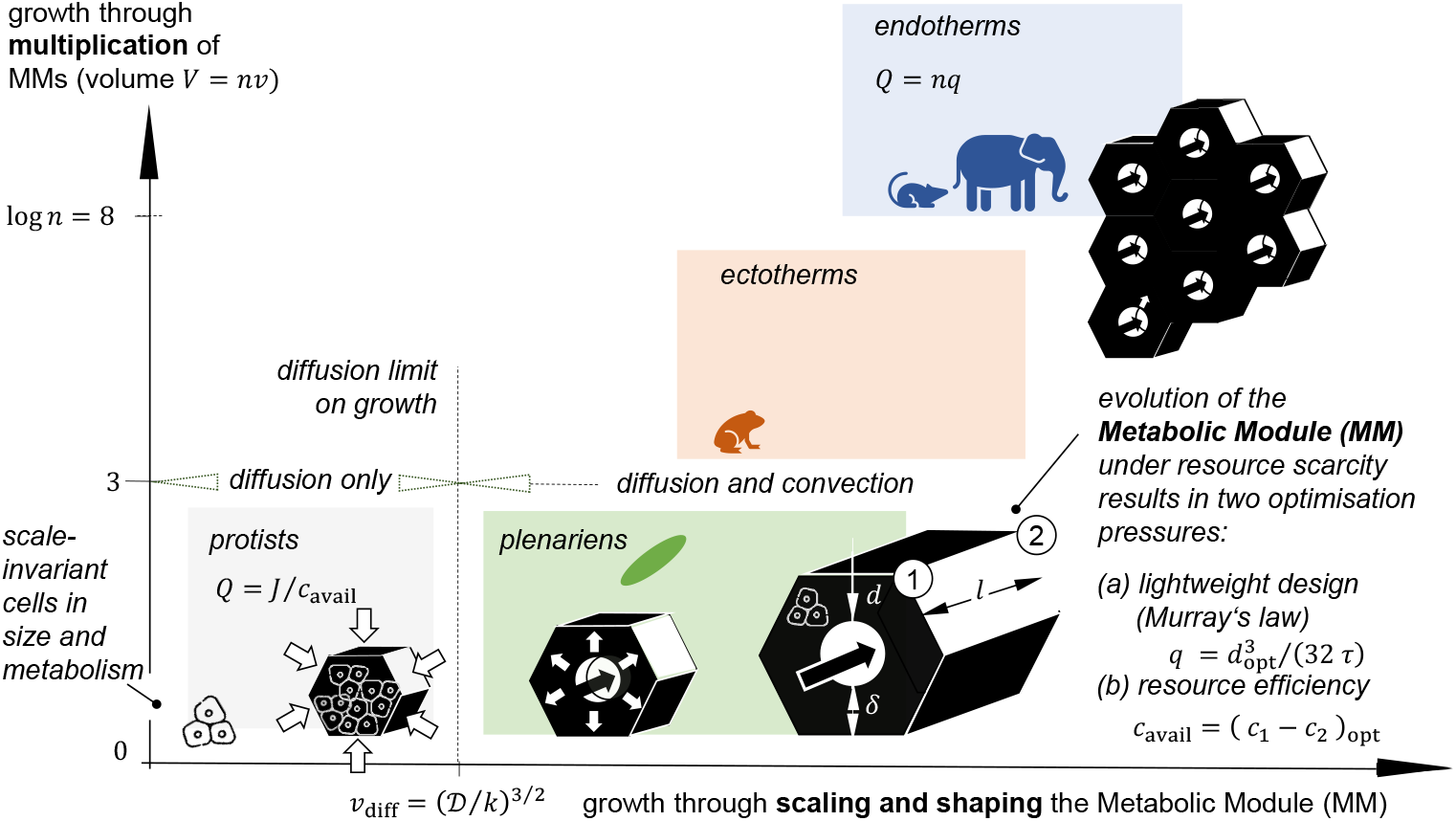
Growth of the organism through first scaling and shaping of the Metabolic Module (MM) and second growth through multiplication of the MMs

The capillary module is a known concept [5]. What is new, however, is that the shape, size and thus also the metabolism of MMs are optimised for the size of the organism through natural selection. Understanding the MM is at the heart of this article, i.e. understanding the scaling of the size and shape of the slenderness of the MM through Darwinian evolution by selection. In fact, this is the key to linking cell metabolism and the metabolism of organisms.

### Consecutive growth strategies

In the new perspective presented here, nature applies two sequential evolutionary growth strategies, which are sketched as vertical and horizontal directions in fig. 1. The horizontal direction shows volume growth by scaling and shaping the MM. The vertical direction shows consecutive growth by multiplication of MMs. For cell tissues and organisms that are smaller than the diffusion limit on growth, growth is achieved by cell multiplication. For organisms above that size limit, the MMs are the optimised modules of the organism, which are scaled and shaped in the first evolutionary step and multiplied by a factor *n≥* 1 in the second evolutionary step.

As a result of the two consecutive growth strategies, the volume *V* = *nν* of the organism is derived from the volume *v* of the module, and the blood volume flow *Q* of the organism is derived from the volume flow *q* through the capillaries of the module as *Q* = *n q*. To obtain the metabolic rate of the organism, measured as heat production per unit time and its mass, both values must be multiplied by the energy and mass density, respectively: *Ė* = *e Q* = *en q, m* = *ϱ V* = *ϱn ν*.

Figure 1 shows that the MM is a cylindrical tissue structure of length *l*, which is supplied with nutrients and oxygen by a concentric capillary of diameter *d*. The tissue thickness is *δ* and the effective outer diameter is correspondingly *d* + 2*δ*. The nutrients and oxygen are first transported by convection in the capillary and finally by diffusive transport within the cell tissue in order to meet the needs of cell metabolism.

### Natural selection pressures caused by resource scarcity

The scarcity of resources and the struggle for survival lead to natural selection caused by an optimisation pressure. In fact, there are two optimisation pressures that scale and shape the MM, namely (a) minimum mass of the MM for a given power (lightweight design) and (b) optimal use of available nutrients and oxygen (resource efficiency). Both are achieved through natural selection, which is simulated here using model-based optimisation.

The principle of lightweight design is also known as the ‘physiological principle of minimum work’ [8]. Lightweight design is essential for high acceleration in animal locomotion. But it is also crucial for two other reasons: firstly, the mass-to-power ratio should be minimised when energy reserves are limited, e.g. during the incubation phase of an egg; secondly, a minimum power-related mass is required to limit the energy expenditure for cell renewal.

a. **Selection pressure for lightweight design** With minimal blood mass and minimal dissipation, the optimal ratio between the blood volume flow *q* through the capillary and its diameter *d* is given by Murray’s law [6, 8] 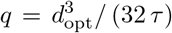. It results from a constrained optimisation problem. The design variable is the capillary diameter *d*, the constraint is the given volume flow *q*, and the objective function to be minimised is the sum of the blood mass and the energy loss due to capillary flow. The given form of Murray’s law with the physiological constant *τ* in the denominator is new; it shows how nature realises Murray’s law: adaptive growth is often triggered by the prevailing shear stress. If Murray’s law applies, the blood cells in capillary flow experience the same wall shear stress and the same wall shear rate, 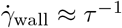, regardless of the capillary diameter, the blood volume flow through the capillary, and the size of the organism and its taxonomic classification. Murray’s law therefore provides an equilibrium state of growth in a cybernetic loop. Like any other closed control loop, it contains a measuring element, a controller and a controlled variable. The latter is the capillary diameter. The measuring elements are the cells forming the capillary wall and measuring wall shear rate. If 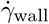 deviates from *τ*^−1^ in a way that is unacceptable for the organism, the organism responds with a change in capillary diameter. This adaptation happens at the cellular level until the wall shear rate is approximately equal to*τ*^−1^ [9–12]. As shown in the section on model validation, *τ*^−1^ is a physiological parameter for blood flow that is known in connection with primary haemostasis. For this reason, too, it can be concluded that Murray’s law forms a solid basis for the theory presented here.
b. **Selection pressure for resource efficiency** In the following, the convective-diffusive transport in capillary and tissue as well as the cell metabolism is considered. Let us call the oxygen concentration at the inlet of the MM *c*_1_ = *c*_basal_ + *c*_avail_ and *c*_2_ at the outlet of the capillary. The capillary is too short for *c*_2_ ≈ *c*_1_, as the allocated and hence available oxygen concentration *c*_avail_ is obviously not utilised. Conversely, the capillary is too long for *c*_2_ ≈ 0, as there is then a risk of cell death in the exit area of the MM. The resources are only utilised efficiently if *c*_2_ assumes the basal concentration *c*_basal_. Resource efficiency therefore leads to (*c*_1_ − *c*_2_)_opt_ = *c*_avail_.

### Mass balances

With the two optimisation results for (a) lightweight design, i.e. 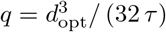, and (b) resource efficiency, i.e. *c*_avail_ = (*c*_1_ −*c*_2_)_opt_, the mass balances for (i) the capillary, (ii) the blood-tissue interface, and (iii) the tissue itself can be formulated.

For this purpose, the integral form of the mass balance is used to develop a concise scaling theory. Alternatively, the corresponding conservation equations can be solved in differential form as a boundary value problem. However, this has no influence on the essence of the theory, as discussed in the outlook.

i. The mass balance for the capillary for resource efficient operation with *c*_avail_ = (*c*_1_ − *c*_2_)_opt_ gives the optimal convective molar flow rate:

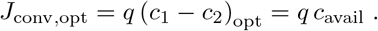
ii. At the blood-tissue interface *S* = *π d l* with the radial normal vector 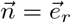, the transport of molecules must be continuous, i.e. *J*_conv,opt_ = *J*_diff_. The order of magnitude of the diffusive molar flow rate is

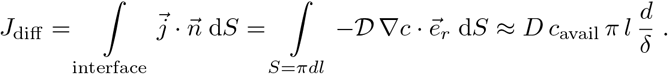 Fick’s first law of diffusion is applied here, in which the molar diffusion flux vector is proportional to the concentration gradient: 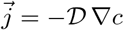. The radial component of the concentration gradient 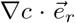 is inhomogeneous. It has a maximum at the capillary inlet and a minimum at the outlet. The right-hand side of the upper equation is therefore an average value for the MM. This is captured by the dispersion coefficient of the cell tissue *D*. This is a parameter that must be much smaller than the diffusion coefficient *D/ 𝒟 ≪* 1 [13, 14]. The factor *D/ 𝒟* can be determined either by solving the reaction-diffusion equation for the cell tissue, as explained in the outlook section of this article, or by using measured metabolic rates. The latter is done in the following section.
iii. The model is closed with the cell metabolism. At each point in the cell tissue, the temporal change in concentration *ċ* is balanced by the diffusive supply of oxygen. The integral form of the reaction-diffusion equation for the volume of the cell tissue *ν*_tiss_ is the same as for ‘small’ animals of volume *ν*, eq. (1), i.e.

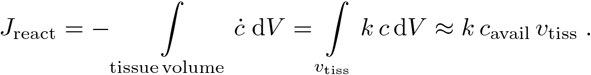

However, the volume of cell tissue in a ‘large’ organism is only a fraction *ν*_tiss_*/ ν <* 1 of the total volume of the MM. In other words: the porosity 1 −*ν*_tiss_*/ ν* of an organism is generally not zero, whereas in ‘small’ organisms, porosity is zero due to the absence of capillaries.

### Kinematic essence of the general model

Using *J* = *q c*_avail_, the geometric relationships, the mass balances *J* = *J*_conv,opt_ = *J*_diff_ = *J*_react_ and Murray’s law, each formulated for a MM, the following five nonlinear equations result:

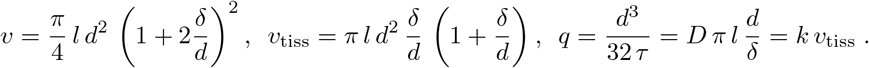

For a given volume flow *q*, the five unknowns *ν, ν*_tiss_, *d, l, δ* are derived from the system of equations - very easily even with a spreadsheet. The solution is the general model for the metabolism of a metabolic module:

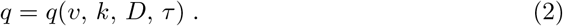

Here, as in the metabolism of ‘small’ organisms, eq. (1), the core of the model, is kinematic in nature, as it represents the optimal transport of molecules within the MM. From a logistical point of view, the MM is designed for a ‘robust just-in-time oxygen supply chain’. The MM is robust because the cell can continue to perform its function of energy conversion for a short time even if the supply chain is insufficient. This robustness is achieved by storing anaerobic energy resources in the cells. The transition from a passive to an active organism in aerobic cell metabolism is discussed in a separate section below. As we will see, the general model, eq. (2), covers the metabolism of both ‘small’ and ‘large’ organisms of different taxonomic classes.

Darwin’s principle of selection for (a) lightweight design and (b) resource efficient solutions was explained above by first modelling the physiology of the MM axiomatically and then optimising the MM based on the model. The central idea is that nature scales and shapes MMs under scarcity using selection heuristics, while model-based optimisation simulates this.

If the concept is viable, and this is demonstrated in the following (i) by model validation using measured metabolic rates, (ii) by independent derivation via dimensional analysis, and (iii) by successful model use for predicting the heart rate of mammals, then both strategies lead to the same functional relationship

## 3 General model - properties and validation

With the metabolic model for one MM, eq. (2), the metabolism of an organism is obtained by multiplying the modules up to the number *n*: *Ė*= *ne q* and *m* = *nϱ ν*.

The only difference between the taxonomy classes protists, planarians, ectotherms and endotherms in the presented theory, see fig. 1, is the number *n* of MMs. The order of magnitude of MMs in mammals is one billion [2], which gives the limits for the number of MMs: 1≤ *n <* 10^9^.

Figure 2 therefore shows the solution of the system of equations as black solid lines for log *n* = 0, 3, 8 as a parameter to distinct the set of curves. The number of MMs for the various taxonomic classes was determined based on measurement data, see fig. 3. The data obtained is consistent with the upper limit provided [2].

**Fig. 2.**
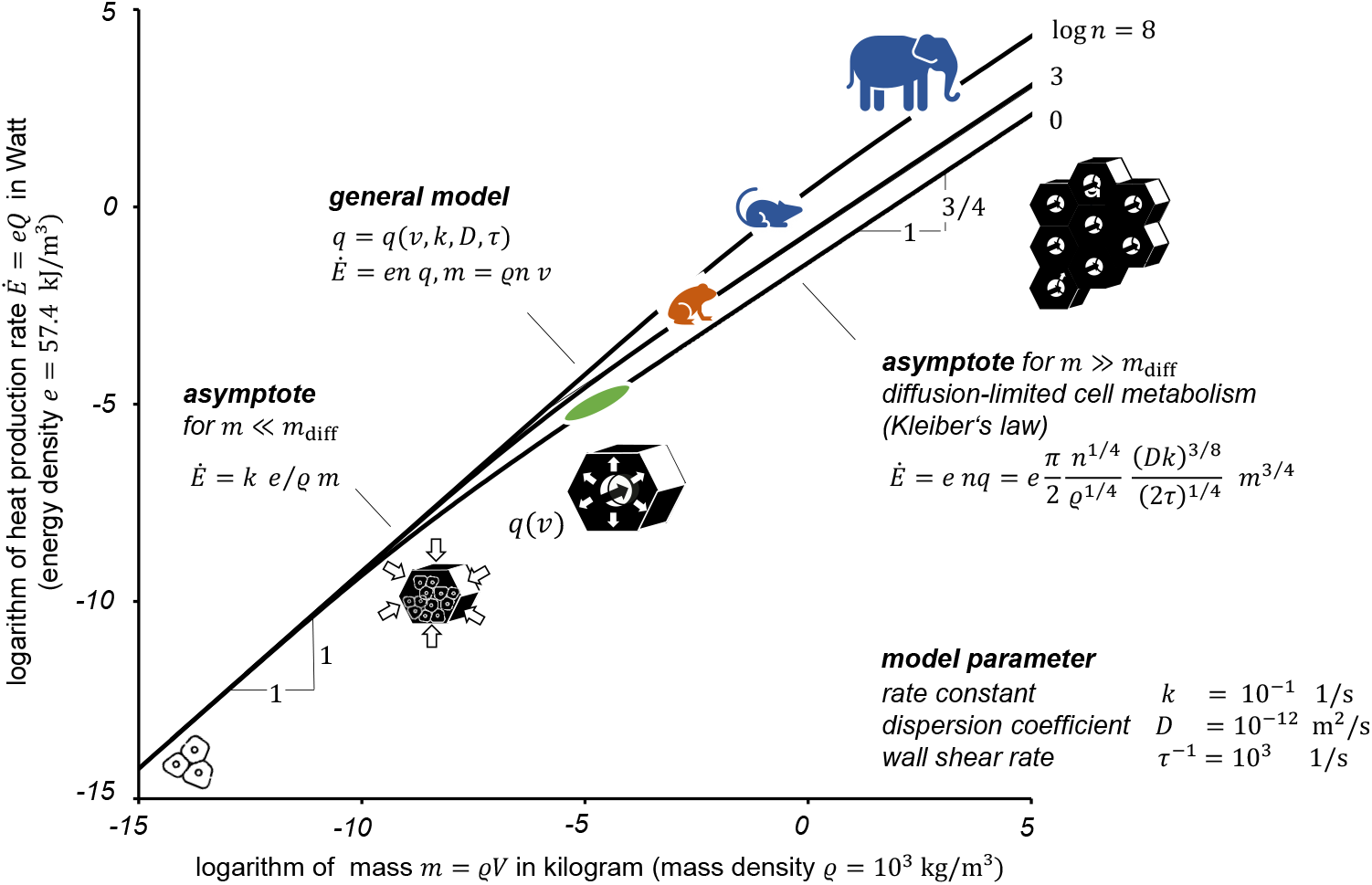
Metabolic rate as a function of organism size for three different numbers of metabolic modules represented by the given theory; all given physicochemical and physiological properties for cell tissue and capillary flow are held constant and assume typical values. The model proves to be a general theory for the entire mass range. The asymptotes for ‘small’ and ‘large’ organisms are clearly visible. In the latter case, the asymptote is due to diffusion-limited cell metabolism.

**Fig. 3.**
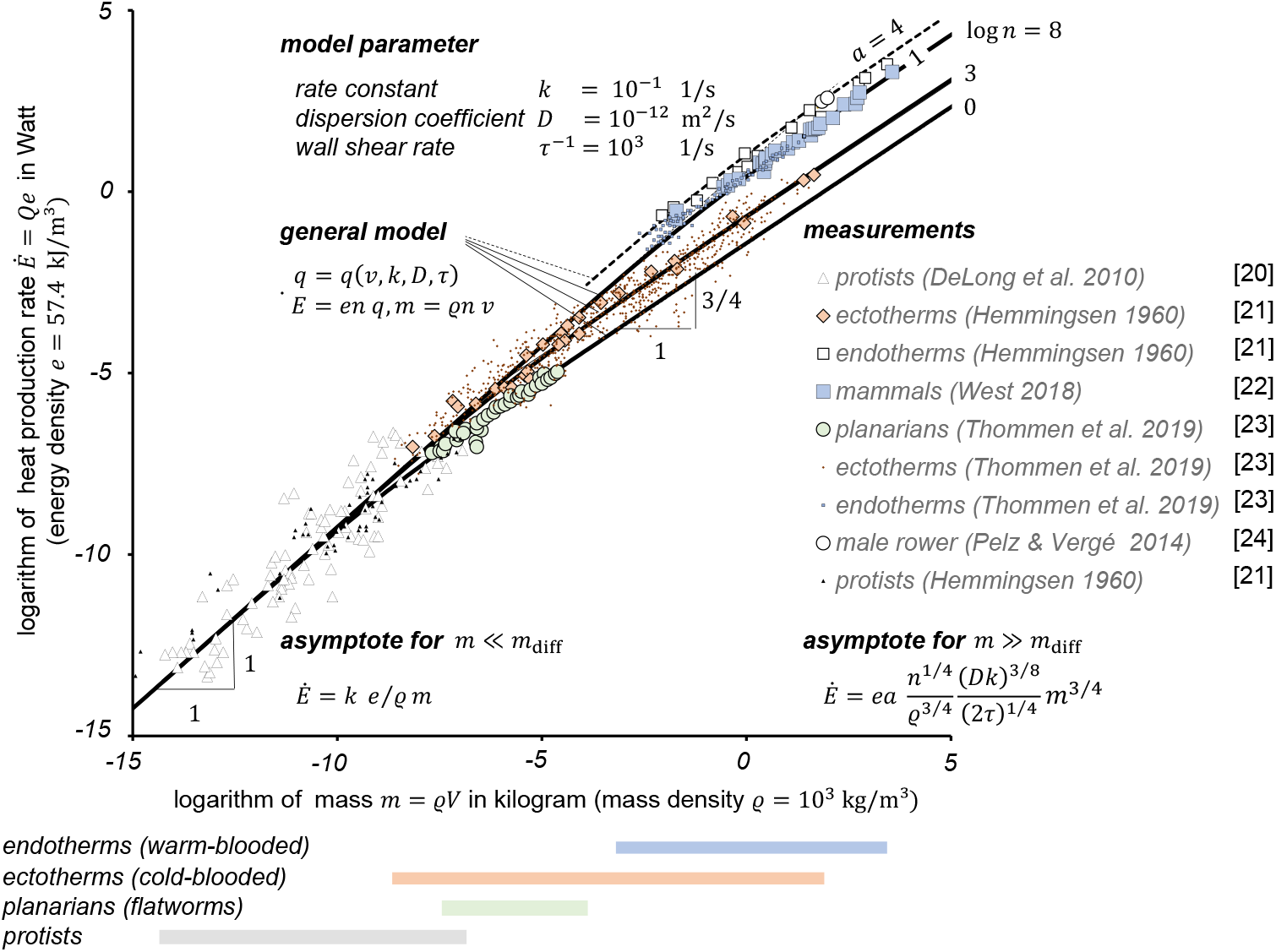
Model vs. measurement: the solid lines show the calculated metabolic rate as a function of organism size for three different numbers of MMs represented by the given theory; all physicochemical and physiological properties for cell tissue and capillary flow are kept constant. The symbols show measured metabolic rates for protists, planarians, ectotherms and endotherms. The dashed line applies to log *n* = 8 and an activity factor of *a* = 4, which corresponds to an increase of heart rate by approximately the same factor. Data are taken from [20–24].

All other physicochemical properties, i.e. *ϱ, e, k, D, τ*, are determined by the tissue and blood physiology and are therefore the same for all organisms, i.e. they are scale-invariant. The following typical parameters are identified on the basis of measurements and experience and used as constant model parameters in the following: The rate constant for glucose oxidation in the temperature range from 25 ^*°*^C to 37 ^*°*^C is *k ≈*0.1 s^−1^ [15]. The change in molar Gibbs free energy for glucose oxidation is Δ*G*^*°*^ =− 2.87 MJ mol^−1^ [16] and the assumed available oxygen concentration is *c*_avail_ = 0.02 mol m^−3^, resulting in the energy density *e* = *c*_avail_ | Δ*G*^*°*^| = 57.4 kJ m^−3^. To prevent primary haemostasis, the required shear rate within the capillaries is *τ*^−1^≈ 10^3^ s^−1^ [9, 10]. The molecular coefficient of diffusion in dilute solutions is *𝒟* = 10^−9^ m^2^ s^−1^ [14]. A comparison of the model results shown in fig. 2 with the measured metabolic rates (see fig. 3) shows that the dispersion coefficient is three orders of magnitude smaller, *D* = 10^−3^ 𝒟. As already mentioned, this takes into account the inhomogeneous concentration gradient field in the cell tissue of the MM. *D* is therefore an effective value for the MM.

Given the conciseness of the model, it is remarkable that the metabolic rate of all taxonomic classes and all organism sizes can be predicted (completeness) not only relatively but also absolutely within the usual model uncertainty of a scaling theory. In addition, three further arguments for the validity of the theory are presented below, namely its asymptotes, its independent confirmation by dimensional analysis, and the application of the model to scale the heart rates of mammals.

## 4 Kleiber’s law is an asymptote of the general model

Figure 2 shows the metabolic rate as a function of body mass. This curve is also known as ‘mouse-to-elephant curve’. Two expected asymptotes *Ė∝ m* and *Ė*∝ *m* ^3/4^ are clearly visible.

Both special solutions are asymptotes of the given system of five nonlinear equations and thus of the general model given by eq. (2): for *δ/d ≫* 1 the asymptotic solution is *q* = *k ν*. For *δ/d ≪* 1, the diffusion limit on growth is reached, and the tissue thickness is limited by the diffusion length *δ →δ*_diff_ = (𝒟*/k*)^1*/*2^. The explicit form of this asymptotic solution is derived from eq. (2) as:

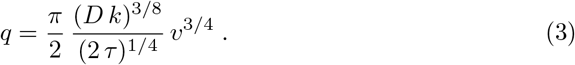

This result is confirmed by a second independent approach using dimensional analysis [17], as shown in the appendix to this article.

For the asymptote, eq. (3), the reaction kinetics in some of the downstream cells are just becoming diffusion-limited. In fact, the MM is optimised in size and shape by natural selection so that the diffusion time is equal to the reaction time, i.e. in logistics terms: in the optimal MM for the asymptotic limit of ‘large’ organisms, the supply chain for oxygen molecules is a just-in-time supply chain. When growing by multiplying a number of *n ≥* 1 MMs with the mass and energy density *ϱ* and *e* respectively, the two asymptotes are given by eq. (1) for *m ≪ m*_diff_ and

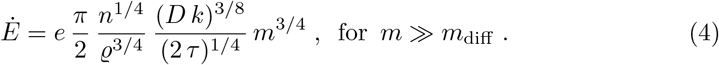

Equation (1) is the asymptote for a metabolism that is not diffusion-limited, eq. (4) is the asymptote for a diffusion-limited metabolism that is supported by optimal design of the organism’s MMs.

In fact, this latter asymptote is Kleiber’s law, which is derived here axiomatically in absolute quantities and not only as a relative scaling law. To the author’s knowledge this is the first link from cell metabolism to the metabolism of the organism. The theory is convincingly confirmed by available experiments, see fig. 3.

## 5 From passive to active

As organisms are at rest most of the time, it can be assumed that the MMs are optimised for this operating state by natural selection. Therefore, the calculated volume flow *Q* = *n q* is for the resting state. For mammals, the ratio of heart volume to body volume is geometrically similar, i.e. *α* := *V*_heart_*/V≈* 6 % independent of the body sizes *V* [18]. The blood volume flow is proportional to the product of heart rate *f* and heart volume: *Q* = *f V*_heart_ = *α f V*. With *Q* = *n q* and *V* = *n v* and the general model, eq. (2), the heart rate at rest is therefore proportional to *f ∝ m*^−1/4^. This analytical model result is validated by experiments [19].

A mammal adapts its metabolism to a required activity by increasing the heart rate by an activity factor *a* := *f*_active_*/f*. For example, a rower’s heart rate can increase from a resting rate of *f* = 50 bpm to *f*_active_ = 200 bpm during a race. The activity factor is therefore *a* = 4. This increases the volume flow to *Q*_active_ = *V*_heart_ *f*_active_ = *V*_heart_ *af* = *a Q*. The metabolic rate for the active organism is hence *Ė*_active_ = *aĖ* or log *Ė*_active_ = log *a* + log *Ė*. This constant shift in the vertical direction can be seen in fig. 3 for log (*n* = 8): the metabolic rate of mammals at rest is the reference, see solid line for *a* = 1 in fig. 3. The dashed line is the same line, but shifted vertically by the amount log (*a* = 4).

## 6 Conclusion and outlook

The paper links the physiology of cell tissue with the metabolism of organisms concisely, clearly and consistently with physiology such as Murray’s law. In this way, the theory presented opens up a new and useful perspective on the metabolism of organisms. It can also be used to provide an insight into evolution under resource scarcity. Darwin’s evolution by selection is interpreted as an optimisation heuristic. This heuristic is rationalised by model-based optimisation of an axiomatic model. The optimisation pressures are directed towards (a) lightweight design and (b) resource efficiency. Together, these two pressures lead to Kleiber’s law.

In this paper, the conservation laws are formulated and solved in integral form. For a detailed analysis of the concentration field in the MM, the convection-diffusion-reaction equation must be formulated and solved in differential form for both the capillary and the cell tissue. This associated boundary value problem can be approached numerically or analytically. The analytical solution leads to an eigenvalue problem in which the eigenvalues are determined by the boundary values. For this, the solution must be developed in a Bessel series. The optimisation with regard to (a) lightweight design and (b) resource efficiency can then be carried out on the basis of the numerical or analytical solution, as in the present work. This would lead to more details, but the essence of the results remains unchanged.

## Appendix

The asymptote, eq. (3), of the general model, eq. (2), i.e. Kleiber’s law, is confirmed by an independent derivation using dimensional analysis [17]. Due to Buckingham’s Pi theorem, the functional relationship of the five kinematic variables *q* = fn(*v, k, D, τ*) is equivalent to that of only three dimensionless variables Π_1_ = fn(Π_2_, Π_3_), with

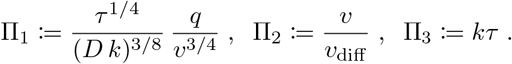

For physical reasons, Π_1_ must be constrained by a constant function value if one of its arguments approaches zero. Therefore, Π_1_ = fn(*ν*_diff_ */ν →*0, *kτ*) const.

The numerical value of this constant, namely *π/*2^5/4^, cannot be derived from dimensional analysis but only by comparison with the analytical result, eq. (3), or by comparison with experiments. Nevertheless, dimensional analysis leads to the same result in a second, independent way, namely *q ∝v*^3/4^, valid for ‘large’ organisms, i.e. for diffusion-limited cell metabolism. In addition to validating the model through experiments, see fig. 3, this independent derivation provides strong support for the theory presented.

## Acknowledgements

I thank the anonymous reviewers for their curiosity and efforts. I would like to thank my co-workers for their supportive team spirit. Finally, I would like to thank my family from the core of my heart for their love, without which life is not possible.

## Declarations

The author declares that he has no competing interests.

